# An exact version of Hunt’s ancestor-descendant directional random walk model

**DOI:** 10.64898/2026.08.21.746177

**Authors:** Rolf Ergon

## Abstract

Hunt’s ancestor-descendant (AD) method for fitting evolutionary models to empirical paleontological sequences assumes independent log-likelihoods for the transitions between populations (Hunt, 2006). This is not quite correct, as he also pointed out in his paper. The reason is that trait differences in adjacent transitions are correlated, and ignorance of this fact may give large errors in the estimated parameter and Akaike Information Criterion (AIC) values. Here, the problem is solved by use of the *N* − 1 – dimensional normal density for a random vector, where *N* is the number of samples. The exact AD model has a tridiagonal trait covariance matrix instead of Hunt’s diagonal matrix, and the estimated prediction slopes in cases where the estimated random component is zero will then be identical to those found by weighted least squares estimation.

## 1. Introduction

Hunt (2008, 2024) presents two different methods for fitting evolutionary models to empirical paleontological sequences. The first uses the mean phenotypic differences between ancestor – descendant (AD) pairs of populations, while the second considers the distribution of all sample means jointly (JOINT), and each of the two models have their weaknesses. As developed under the name general random walk (GRW) model (Hunt, 2006), the original AD model makes use of the log-likelihood for each single ancestor – descendant transition, i.e., for *N* − 1 transitions between *N* samples. The total log-likelihood is then found by summing the log-likelihoods for all transitions, and that introduces an error caused by the “dependence between adjacent trait differences because they share a sample mean and its sampling error” (Hunt, 2006). The problem was illustrated in Ergon (2026), where the random evolutionary components in four real data cases were found to be zero. Although Hunt’s AD prediction slope parameters in these cases were found by maximum likelihood estimation, they were different from optimal weighted least squares (WLS) results, which indicates that the Hunt’s AD model is incorrect. Some of these cases show that the error in Hunt’s AD model may have substantial consequences. The main aim of this article is to find a solution to this problem. The JOINT approach, on the other hand, tends to underestimate the variance of the random walk component, which leads to an increased risk of obtaining negative step variances.

The known theoretical background and the solution to the AD problem is given in Section 2, where also new results for cases with zero random components are included. The results are illustrated by simulations in Section 3 and by real data cases in Section 4. A summary with discussion and conclusions follows in Section 5.

## 2. Methods

### 4.4 Hunt’s AD model

As developed by Hunt (2006) under the name general random walk (GRW), Hunt’s AD model is based on the log-likelihood function for a single evolutionary ancestor – descendant (AD) mean trait transition,

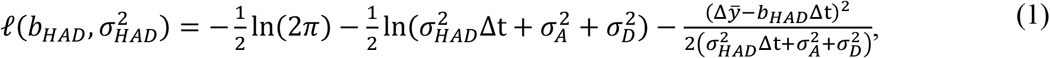

where *b*_*HAD*_ and 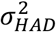 are the mean value and variance of normally distributed incremental mean trait steps, while 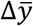 and Δt are the change in mean trait value and elapsed time, respectively. Here, 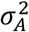 and 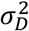 are the phenotypic variances for ancestor and descendant populations, respectively. Note that Hunt (2006) used the notations *b*_*HAD*_ = *μ*_*step*_, 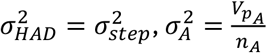 and 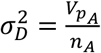. With *N* sparse samples of mean trait values, multiple ancestor-descendant mean trait differences over *N* − 1 evolutionary transitions may according to Hunt (2006) be used jointly to estimate *b*_*HAD*_ and 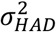 by summing the log-likelihoods according to Eq. (1) over the transitions, i.e., by use of

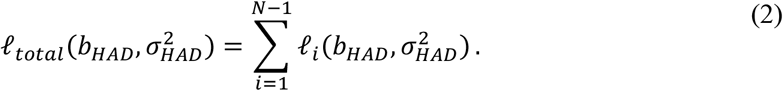

The step parameters in Eq. (2) can be found by maximizing 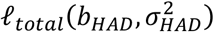, i.e., by maximum likelihood estimation, and the estimated step size 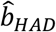 will then be a measure of directional change over time. From the estimated parameter values also follow AIC values needed for model selection (Hunt, 2006). As discussed in Hunt (2006), there is an error in Eq. (2) caused by the “dependence between adjacent trait differences because they share a sample mean and its sampling error”, and a solution to this problem follows in Subsection 2.3 below.

### 2.2 The JOINT model

The JOINT model makes use of the *N* – dimensional normal density for a random vector (Ch. 4, Johnson and Wichern, 2008), which in this setting gives the probability density function

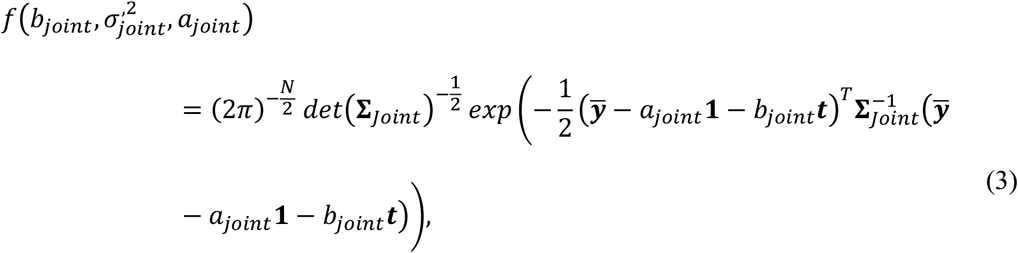

from which follows the log-likelihood function

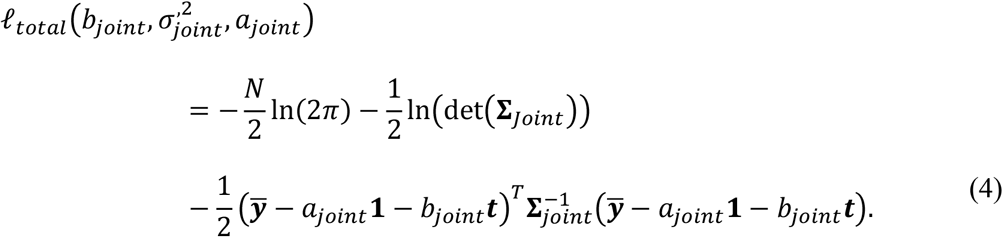

Here, 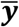 and ***t*** are the data vectors, while **1** is a vector of ones. The trait covariance matrix is 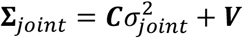, where *c*_*i,j*_ = min(*t*(*i*), *t*(*j*)), while ***V*** is a diagonal matrix of measurement variances (see Ergon (2026) for details of ***C***). Note that a third parameter *a* also must be estimated for alignment of the prediction line *b*_*joint*_*t* with the data in an optimal way.

### 2.3 Solution to the AD parametrization problem

For an exact solution to the AD problem, we must use the *N* − 1 – dimensional normal density for a random vector, which in the AD setting gives the probability density function

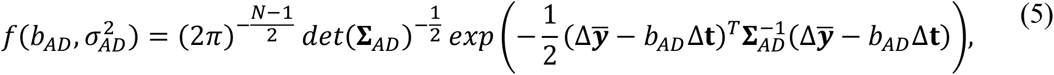

where 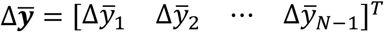 and Δ**t** = [Δt_1_ Δt_2_ ⋯ Δt_*N*−1_]^*T*^. From Eq. (5) follows the log-likelihood function

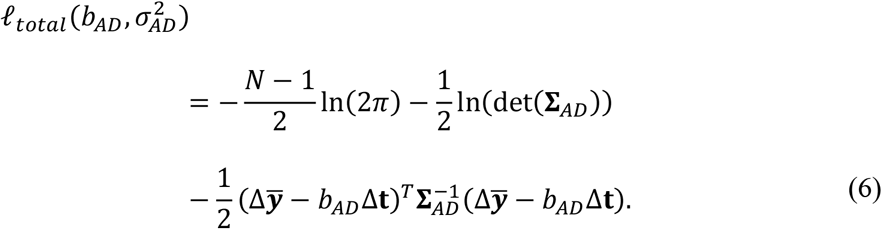

Here, we may for simplicity assume *N* − 1 = 9 transitions and find the tridiagonal band matrix

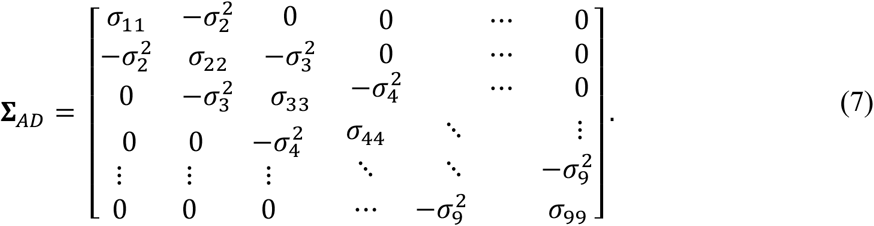

Noting that individual measurements are 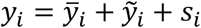, where 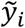 is the measurement error while *s*_*i*_ is the random walk component, we find the diagonal elements

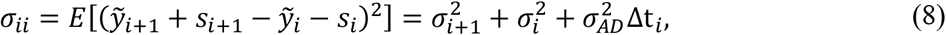

where 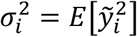 while 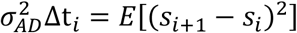 is the increase in variance of the random walk component over the time interval Δt_*i*_. We also find the off-diagonal elements

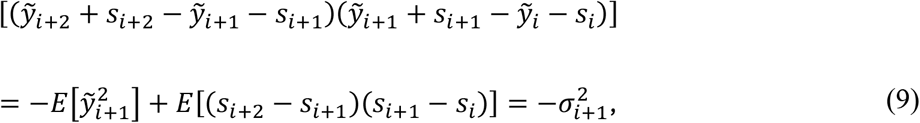

where *E*[(*s*_*i*+2_ − *s*_*i*+1_)(*s*_*i*+1_ − *s*_*i*_)] is zero because *s*_*i*+2_ − *s*_*i*+1_ and *s*_*i*+1_ − *s*_*i*_ are independent. Note that ignoring the off-diagonal elements in **Σ**_*AD*_ by setting them to zero will give the same results from Eq. (6) as from Eq. (2).

### 2.4 Weighted least squares estimation

In many simulation realizations and in all three real data cases in Section 4 the realistic step variances are found to be zero, 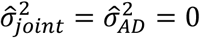, such that the trait covariance matrix is ***V***, where ***V*** is the diagonal matrix of measurement variances. In such cases 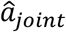 and 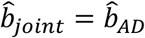 can be found by weighted least squares (WLS) estimation, i.e., from

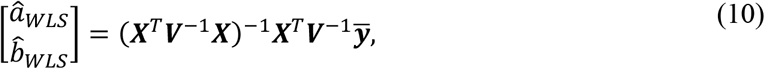

where ***X*** = [**1 *t***]. For the special case with 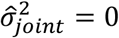 we may now formulate the following theorem:

#### Theorem 1

For 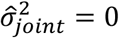 the maximum likelihood estimates *â*_*joint*_ and 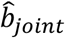 following from Eq. (4) are identical to the WLS estimates found from Eq. (10).

**Proof:** Set [*a*_*joint*_ *b*_*joint*_]^*T*^ = ***θ***. With 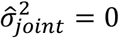 we find **Σ**_*joint*_ = ***V***, and from Eq. (4) we thus find the parameters that maximize 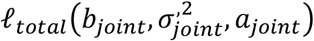 by setting

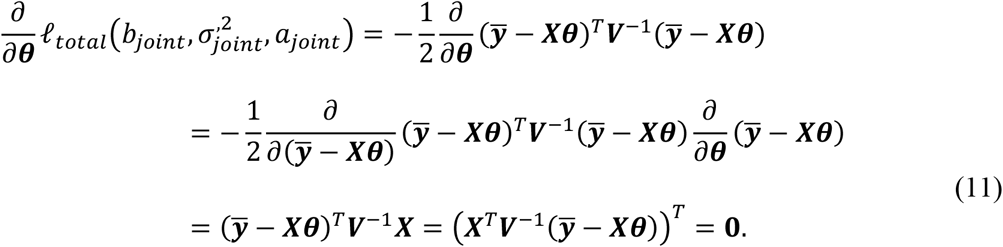

This results in

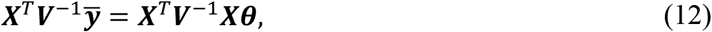

i.e., 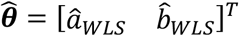 as in Eq. (10).

For 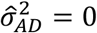 we may in addition formulate the following theorem:

#### Theorem 2

For 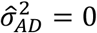 the maximum likelihood estimate 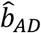 following from Eq. (6) is identical to the slope parameter 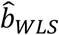 found from Eq. (10).

**Proof** Since the exact AD model with 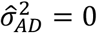 is a purely deterministic maximum likelihood model, and since the data vectors 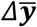 and Δ**t** carry exactly the same information as the data vectors 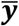 and ***t***, and since in addition the same sample variances are used, it follows that 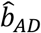 found from Eq. (6) are equal 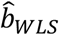 as found from Eq. (10).

## 3. Simulations

### 4.4 Data generation and AD predictions

In the simulations time series data over 1,000 steps of length 1 were generated as

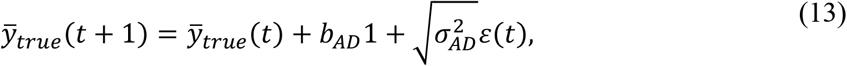

where *ε*(*t*) was a normally distributed random number with variance one. The initial value was chosen to 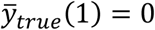, and parameter values were *b*_*AD*_ = 0.1 and 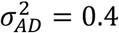 (increased from 0.1 in Ergon, 2026), which over 1,000 timesteps results in an approximately linear change of 100 in 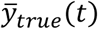. From such time series 10 irregular samples were extracted with *t* as uniformly distributed random numbers in the ranges 1 − 100, 101 − 200, …, 901 − 1000, and to each of these samples was added the mean value of *n* individual measurement values drawn from normally distributed populations with mean zero and variance *V*_*p*_ = 400, resulting in 10 samples 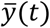. Here, *n* varied between 3 and 60 with uniform random distributions, and the samples thus had standard errors 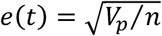 that varied between 2,58 and 11.55.

Based on the generated 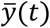 data, estimates 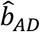 and 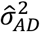 that maximize the exact AD log-likelihood function in Eq. (6) were found. The maximizations were performed using the function *fmincon* in MATLAB, i.e., 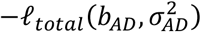 was minimized. The lower limit for the step variance was set to 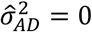.

When 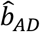 had been determined, a prediction model

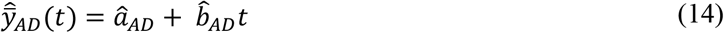

was found by fitting a parameter *a*_*AD*_ to the data by a weighted least squares approach. Such predictions are here used for comparisons in examples of time series responses.

Simulations as described above were also performed using Hunt’s log-likelihood function in Eq. (2), as well as the JOINT log-likelihood function in Eq. (4).

### 3.2 Generalized least squares estimation of reference parameters

In the simulations, maximum likelihood estimates 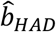 and 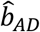 according to Eqs. (2) and (6) were compared with generalized least squares (GLS) estimates 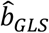 found from

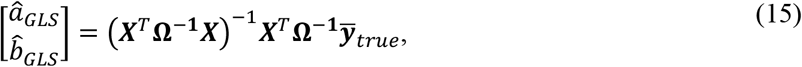

where 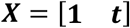 and 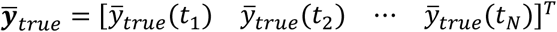, while 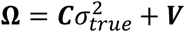, where *c*_*i,j*_ = min(*t*(*i*), *t*(*j*)) (Ergon, 2026). Note that *â*_*GLS*_ and 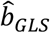 gives the best linear unbiased estimator (BLUE) given the *t* and 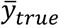 data.

### 3.3 Simulation results with 10 samples

Table 1 shows estimated parameter values for five realizations with *N* = 10 samples and based on the exact AD model in Eq. (6). Results for 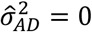 (green numbers) show that 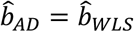, in accordance with Theorem 2.

**Table 1.**
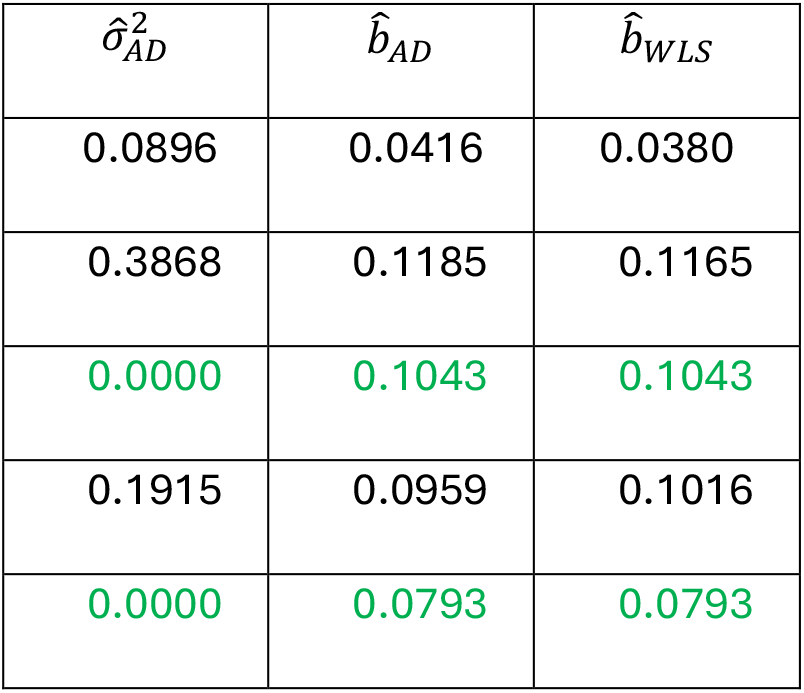
Simulation results for the exact AD model in Eq. (6).

| $\hat{\sigma}_{AD}^2$ | $\hat{b}_{AD}$ | $\hat{b}_{WLS}$ |
| --- | --- | --- |
| 0.0896 | 0.0416 | 0.0380 |
| 0.3868 | 0.1185 | 0.1165 |
| 0.0000 | 0.1043 | 0.1043 |
| 0.1915 | 0.0959 | 0.1016 |
| 0.0000 | 0.0793 | 0.0793 |

Fig. 1 shows histograms using Hunt’s original AD model (Eq. (2), left panels), as compared to use of the exact AD model (Eq. (6), right panels). The upper panels show results for the GLS model in Eq. (11), the middle panels show prediction slope errors for HAD and exact AD models relative to GLS, and the lower panels show prediction slope errors for WLS relative to GLS. The results are based on *N* = 10 samples and 1,000 realizations. Note the differences in the middle panels between the SE values for Hunt’s AD model and the exact AD model. Also note the very similar prediction errors for the exact AD model and WLS. The differences between the two upper panels and between the two lower panels are due to different sets of realizations. Responses for a typical realization are shown in Fig. 2. Here, we can see the well-known effect that the GLS response is very much influenced by the early sample, while 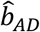 and 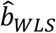 are approximately equal to 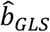. Also note the large error in the prediction slope 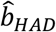 for Hunt’s AD model.

**Figure 1.**
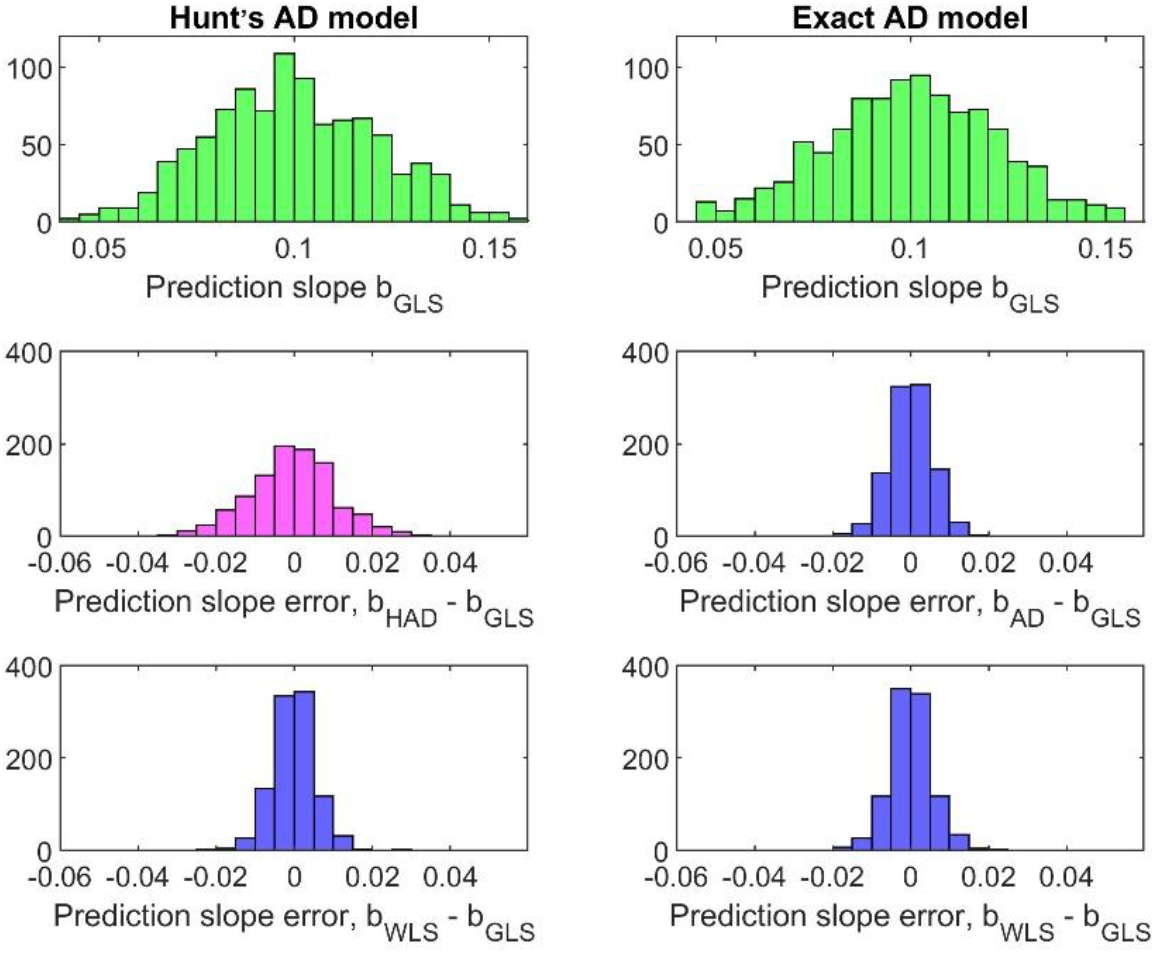
Histograms using Hunt’s original AD method (Eq. (2), left panels), as compared to use of the exact AD method (Eq. (6), right panels).

**Figure 2.**
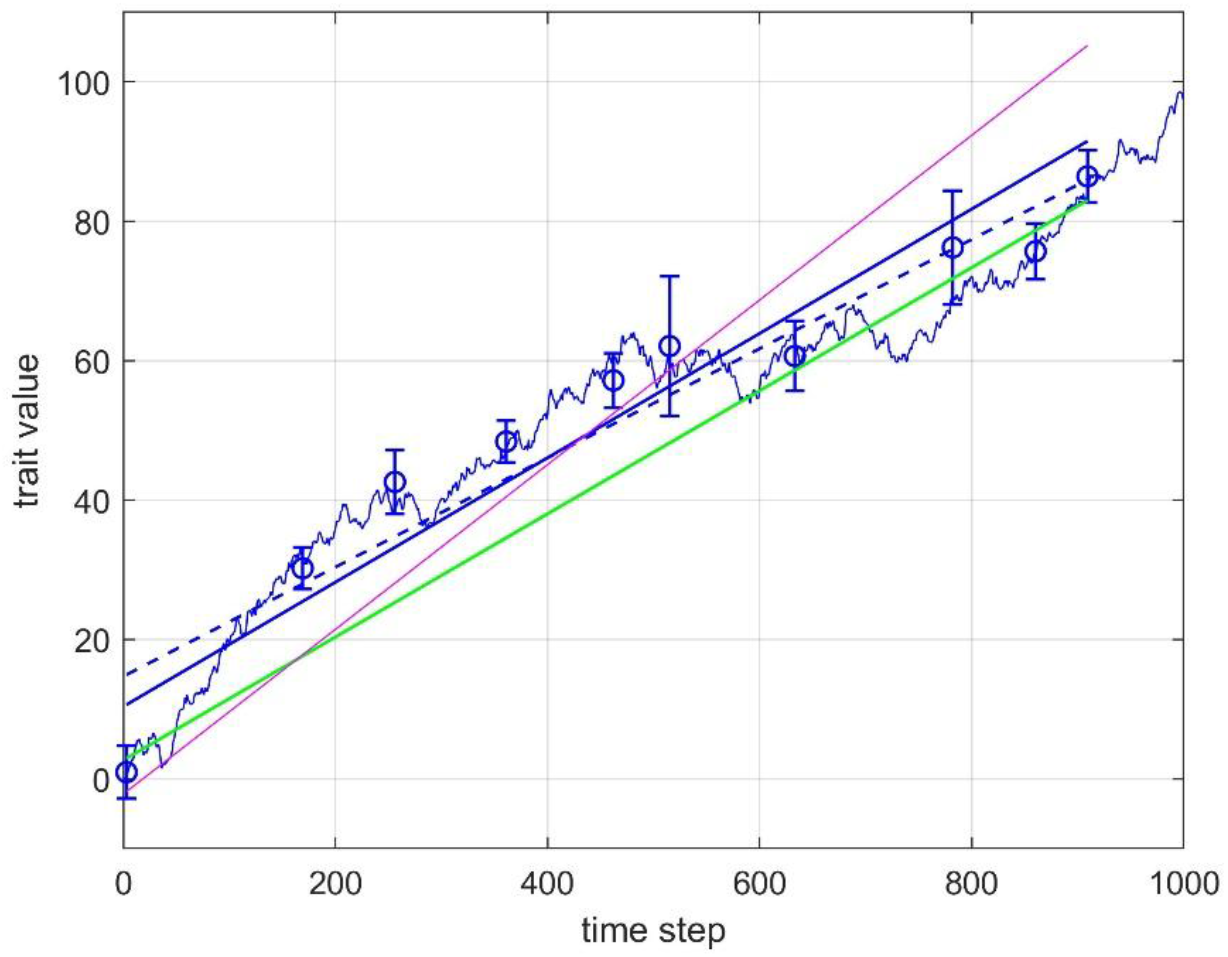
Example results for simulations with 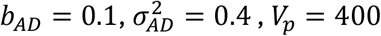 and *N* = 10 samples. Values of 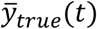 are shown by noisy blue line, while 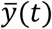 are shown by circles with error bars. GLS predictions are shown by green line. Predictions from the exact AD model are shown by solid blue line, while WLS predictions are shown by dashed blue line. Estimated parameter values were 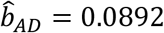 and 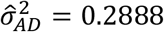. For the same realization of 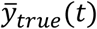, use of Hunt’s AD model resulted in 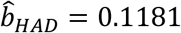 and 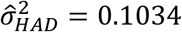, which gave predictions as shown by red line.

### 3.4 Comparison of the AD and JOINT models

Parameter estimates for the exact AD and JOINT models using *N* = 10 samples are given in Table 2. Note that both models tend to underestimate 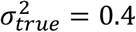, but that this is more pronounced for the JOINT model. This was also noted by Hunt (2008), and it increases the risk of obtaining negative step variances.

**Table 2.** Parameter estimates for the exact AD and JOINT methods, based on *N* = 10 samples and 1,000 realizations, and using the lower search limit 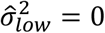. Results are shown as *mean* ± *SE*. All realizations with 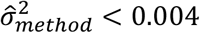 were omitted from the calculations.

| Parameters | Exact AD model | JOINT model |
| --- | --- | --- |
| $\hat{\sigma}_{GRW}^2$ | $0.371 \pm 0.262$ | $0.355 \pm 0.243$ |
| $\hat{b}_{GRW}$ | $0.100 \pm 0.022$ | $0.100 \pm 0.022$ |
| $\hat{b}_{WLS}$ | $0.100 \pm 0.023$ | $0.100 \pm 0.024$ |
| Portion of samples with $\hat{\sigma}_{method}^2 < 0.004$ | 25% | 36% |

Hunt (2024) claims that the JOINT approach is clearly better than his AD approach in cases of long, noisy trends, and simulations were therefore repeated with *N* = 40. Typical responses are shown in Fig. 3, where we again can see the well-known effect that the GLS response is very much influenced by early samples. Results are given in Table 3, which also include AIC values computed as (Hunt, 2006)

**Table 3.**
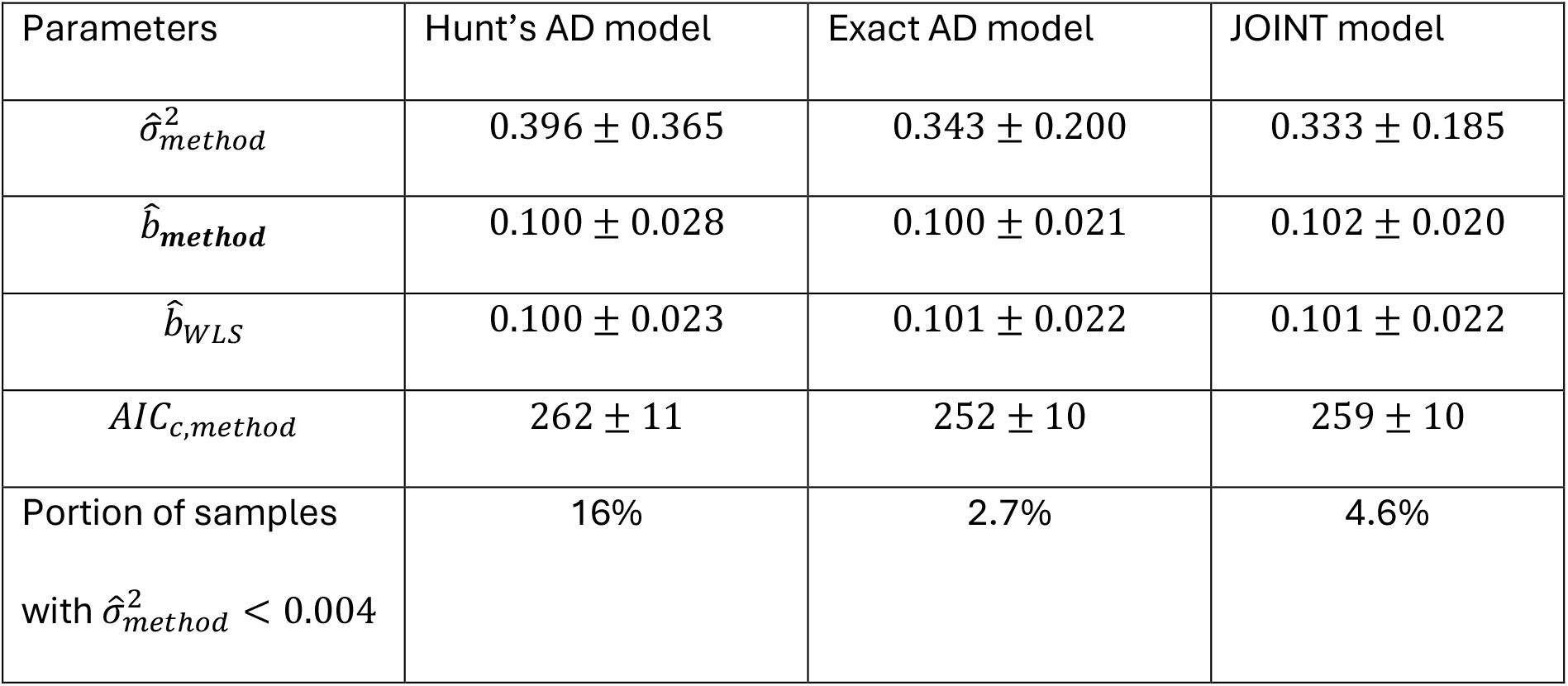
Parameter estimates and AIC values for different models, based on *N* = 40 samples and 1,000 realizations using the lower search limit 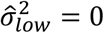. Results are shown as *mean* ± *SE*. All realizations with 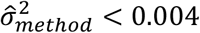 were omitted from the calculations.

**Figure 3.**
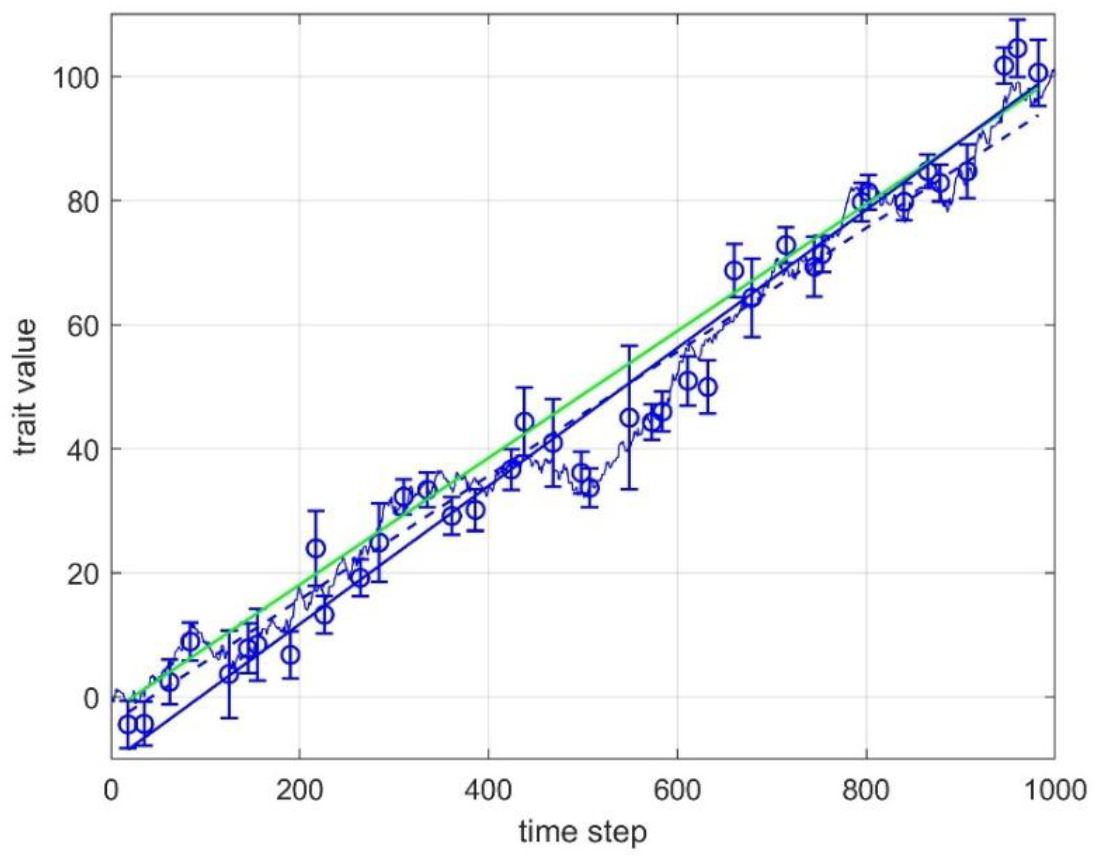
Example results for simulations with *b*_*AD*_ = 0.1, 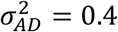, *V*_*p*_ = 400 and *N* = 40 samples. Values of 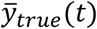 are shown by noisy blue line, while 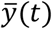 are shown by circles with error bars. GLS predictions are shown by green line. Exact AD predictions are shown by solid blue line, while WLS predictions are shown by dashed blue line. Estimated parameter values were 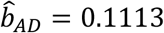 and 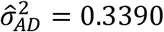.

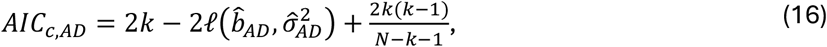

with *k* = 2 and *N* = 39, and

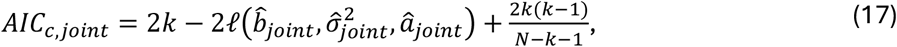

with *k* = 3 and *N* = 40, respectively. All realizations with 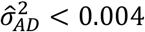 or 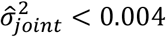 were omitted from the calculations. Note, however, that the AD and JOINT models use different response variables, such that the AIC results cannot be compared for model selection purposes (Burnham and Anderson, 2002). Note that the JOINT model also here tends to underestimate 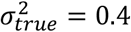 more than the exact AD model does, which results in an increased number of 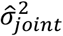 values at the zero limit.

The results in Table 3 confirm Hunt’s claim that the JOINT model performs best of the models now available, in that the number of 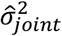 values close to zero is very much reduced, but they also show that the exact AD model performs even better. Although the extra underestimation of 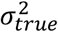 in the JOINT model is less prominent than in Table 2, it is still present, and as a result the number of exact AD realizations with 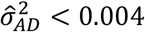 is just over half of the corresponding number for the JOINT model.

Fig. 4 shows four realizations with *b*_*AD*_ = 0.1, 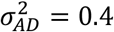, *V*_*p*_ = 400 and *N* = 40 samples, including predictions using both the exact AD and JOINT models, as well as AIC values. Note that the exact AD and JOINT predictions are close to identical, while the AIC values are quite different.

**Figure 4.**
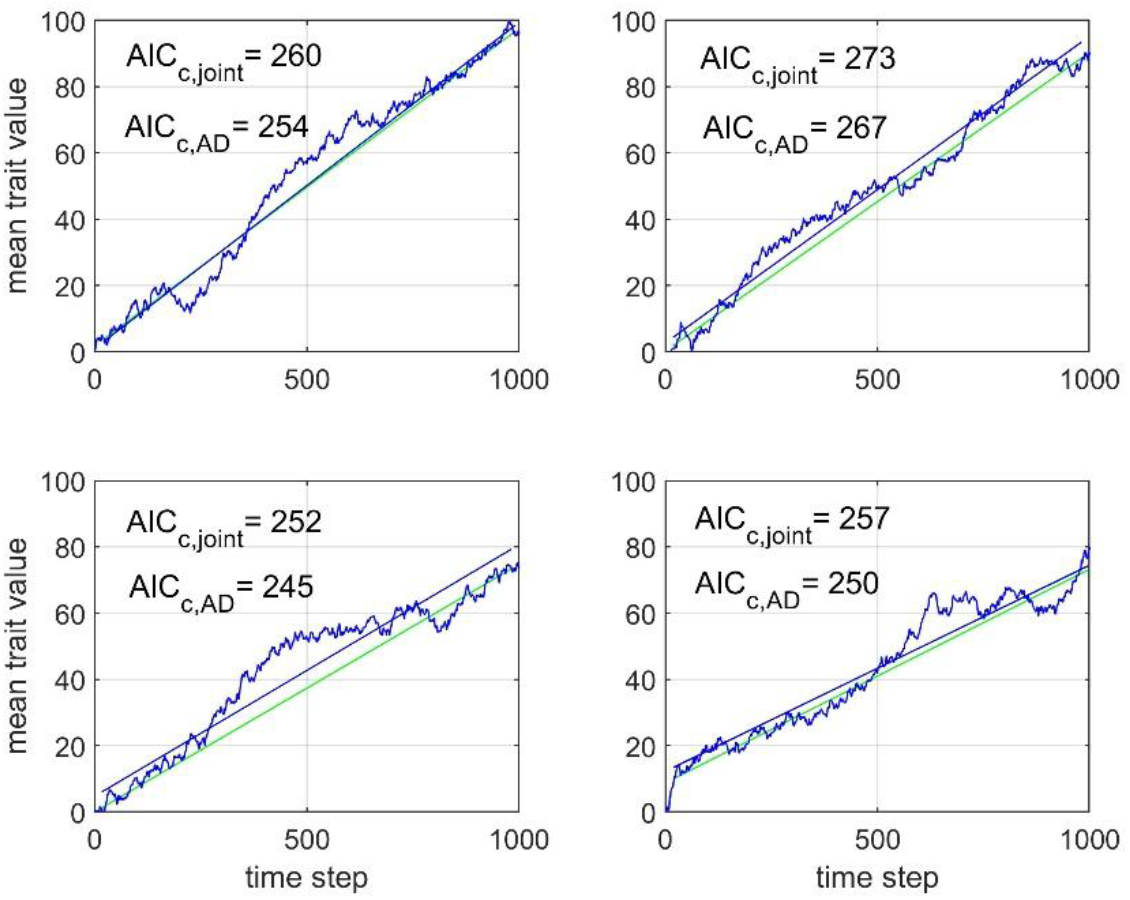
Four realizations using 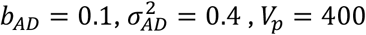 and *N* = 40 samples. Values of 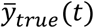 are shown by noisy blue lines, while GLS predictions are shown by green lines. Exact AD and JOINT predictions are shown by close to identical blue lines, and AIC values are given in the plots.

## Real data cases

Fig. 5 shows response plots for the two Ostracod cases in Ergon (2026). Here, the lower limit 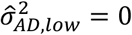 was used in the numerical search, in both cases resulting in 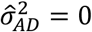. The exact AD model gave 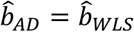 in accordance with Theorem 2, which as shown is quite different from the results from Hunt’s AD model. A closer look at the numerical differences between the prediction slope results in the Ostracod 1 case showed that 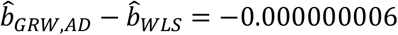, while 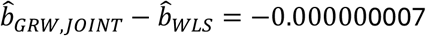.

**Figure 5.**
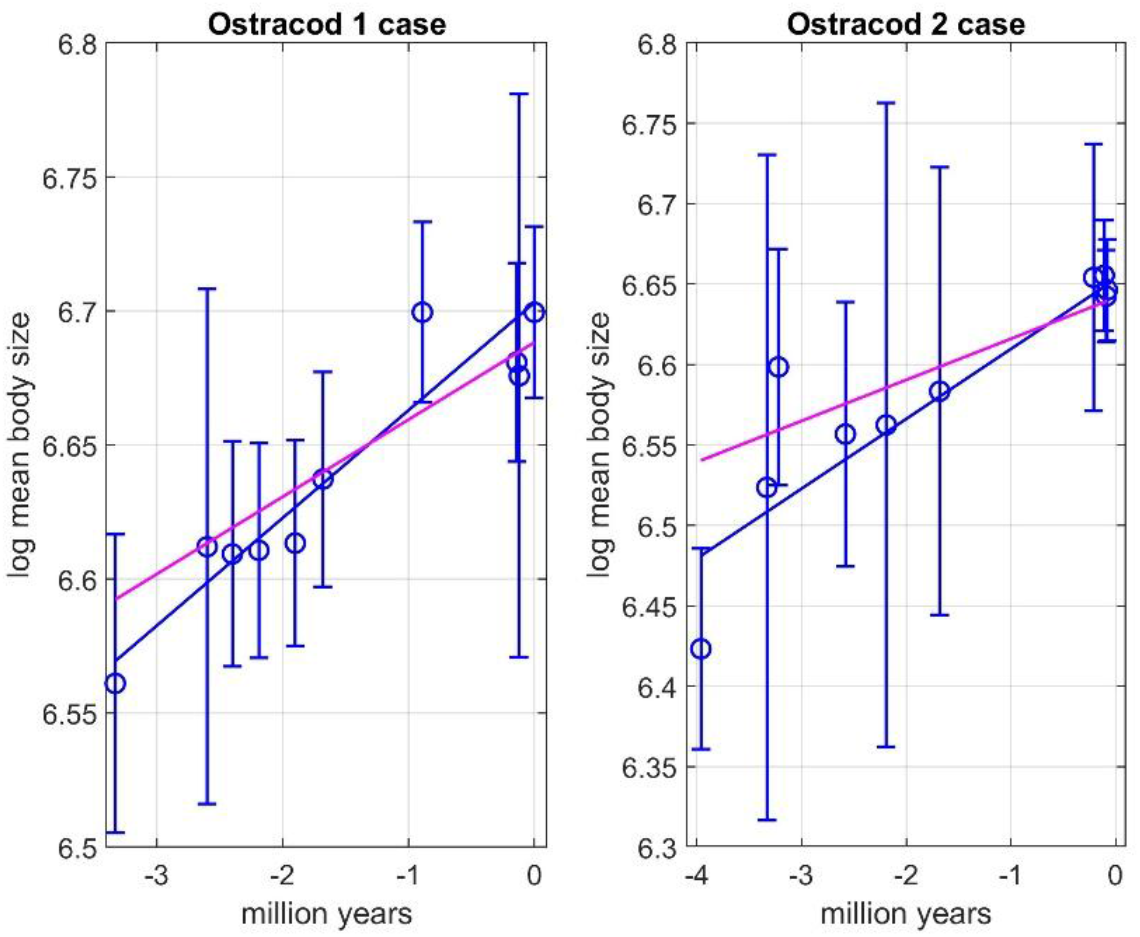
Results for Ostracod case 1 and 2 in Ergon (2026), with mean trait observations (circles with error bars), predictions by use of Hunt’s AD model (red lines), and identical predictions by use of WLS and the exact AD model (blue lines).

The Stickleback fish case in Ergon (2026) was based on data from Bell et al. (1985). The modeling is here modified by using the exact AD and JOINT models, with both log mean traits as in Ergon (2026) and original mean traits as in Bell (1985). The lower step variance bounds were 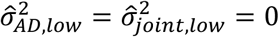, which with log mean traits resulted in 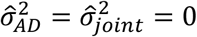. Identical parameter values were then obtained from reduced models with fixed values 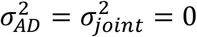. In accordance with Theorems 1 and 2 the estimated parameter values were in these cases identical to the WLS results. Use of the original traits gave positive and very similar step variances and slope parameters. Parameter and AIC results are given in Table 4, and prediction responses based on the original mean traits are shown in Figures 6. Responses based on log mean traits are included in Ergon (2026). Note that the AIC values for the exact AD and JOINT models cannot be used for model selection because the response variables are different.

**Table 4.** Results for the Stickleback fish case.

| Parameter | Exact AD model<br>orig. mean data | Exact AD model<br>log mean data | JOINT model<br>orig. mean data | JOINT model<br>log mean data |
| --- | --- | --- | --- | --- |
| $\hat{\sigma}_{method}^2$ | 5.3395 | 0 | 5.1329 | 0 |
| $\hat{a}_{method}$ | 8.6687 | 2.1465 | 8.6685 | 2.1465 |
| $\hat{b}_{method}$ | 9.5150 | 1.2605 | 9.5187 | 1.2605 |
| $\hat{a}_{WLS}$ | 8.5346 | 2.1465 | 8.5346 | 2.1465 |
| $\hat{b}_{WLS}$ | 11.6424 | 1.2605 | 11.6424 | 1.2605 |
| k | 2 | 1 | 3 | 2 |
| $AIC_c$ | -3.39 | -115 | -0.97 | -121 |

**Figure 6.**
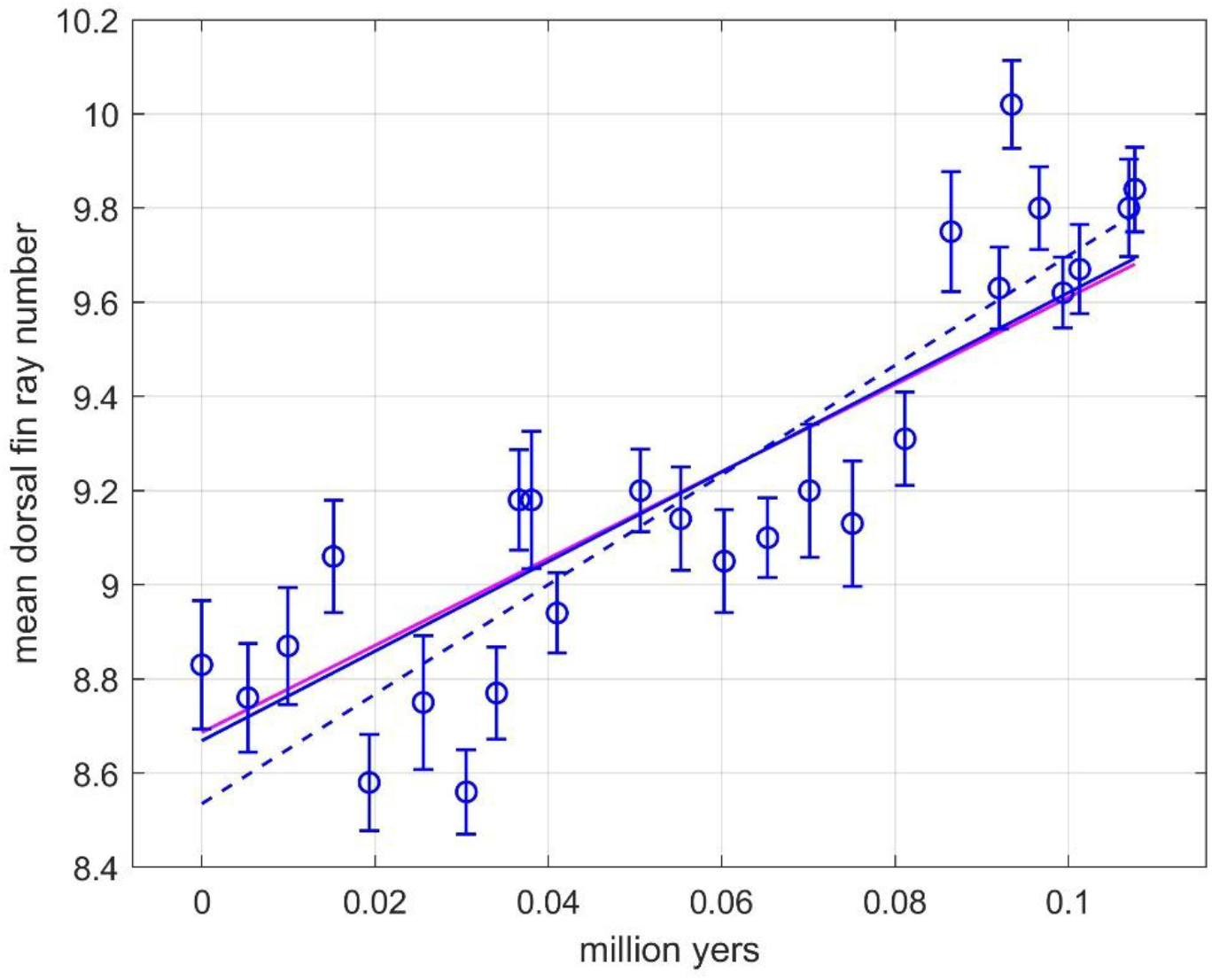
Results for stickleback fish case using original mean trait data, with mean trait observations (circles with error bars), predictions by use of Hunt’s AD model (red line), predictions by use of WLS (dashed blue line), and very similar predictions by use of the exact AD and JOINT models (blue lines). Note that this case results in only a minor difference between Hunt’s AD and exact AD prediction lines.

## 5. Summary and discussion

From a theoretical point of view there are two new results in this article, and the first of these have documented large effects on practical prediction results. Two additional results are of merely practical but not unimportant interest. First, the well-known error in Hunt’s ancestor-descendant (AD) method for fitting evolutionary models to empirical paleontological sequences (Hunt, 2006) is corrected. As shown in Fig. 1, 2 and 5, this may give clearly improved prediction results in cases with realistically large measurement errors. The essential point in this correction is that Hunt’s diagonal covariance matrix in the *N* − 1 – dimensional normal density for a random vector is replaced by a tridiagonal matrix, where the non-diagonal elements account for the correlations between adjacent transitions.

Second, in cases where the estimated step variance is zero, both the exact AD and JOINT models give prediction slope results that are identical to weighted least squares (WLS) results (Theorems 1 and 2, Tables 1 and 4, Fig. 5 and Fig. 7). In such cases both the exact AD and JOINT models will just as WLS result in the best linear unbiased estimator (BLUE).

Third, the results for the real data cases in Section 4 raises an interesting question about estimated slope values for cases where the maximum likelihood point in the parameter space cannot be reached because the step variance has the lower limit 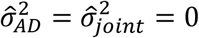. In these cases, the step variances can simply be fixed to zero, and the prediction slope values found by the exact AD and JOINT models will then in accordance with Theorems 1 and 2 be identical to the WLS results. Also note that Theorem 2 assumes an exact AD model, and the difference between 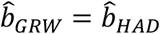 and 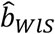 in some of the real data cases in Ergon (2026) were thus clear signs of the error in Hunt’s AD model.

The fourth and practical result follows from simulation tests of the JOINT model, which Hunt (2008, 2024) presents as an alternative to his AD parameterization, and which is now the default choice in the R package paleoTS. The JOINT approach tended to underestimate the step variance, which resulted in an increased risk of obtaining step variances at the lower limit zero (Tables 2 and 3). This step variance underestimation is also seen with use of the original data in the Stickleback case in Section 4 (Table 4).

Simulations are mainly performed using short data with *N* = 10 samples. However, since Hunt (2024) claims that the JOINT approach is better than his AD approach in cases of long, noisy trends, simulations were repeated with *N* = 40. The results in Table 3 confirm Hunt’s claim, but they also show that the exact AD model performs even better, in the sense that the risk for step variances at the lower limit zero is lower.

The situation is thus that the AD parameterization implemented in paleoTS (Hunt, 2024) is theoretically wrong, while the alternative JOINT parameterization gives an increased risk of obtaining zero step variances. This risk is especially pronounced for short data (Table 2), and it will increase for lower values of true step variance. The major result in the present article is that the error in Humt’s AD model is corrected, and the resulting exact AD model performs well. However, whether the exact AD or JOINT models should be used in a wider model comparison context depends on the response variables used in competing models.

The approach behind the exact AD model may have relevance for similar models, for example for the stasis model in Hunt (2006). Details of this are left for further research. With the exact AD model being established, another question for future research is how estimated prediction slopes are affected by unavoidable errors in fossil data sampling times.

## Supporting information

MATAB Code

## Acknowledgments

I thank Gene Hunt and Kjetil Lysne Voje for constructive comments on earlier versions of the manuscript, and University of South-Eastern Norway for support and funding.

## Data Availability Statement

MATLAB code is archived on biorxiv, https://doi.org/10.64898/2026.08.21.746177

