## Supplementary material for "An exact version of Hunt’s ancestor-descendant directional random walk model": MATAB Code

### MATLAB code for

Rolf Ergon

University of South-Eastern Norway

August 20, 2026

#### Figure 1, left panels

```
clear
mu_step=0.1;
var_step=0.4;
Vp=400;
count=0;
y0=0;
N=10;
M=1000

for m=1:M
%% Generate data
y=y0*zeros(1,1000);
for t=2:1000
   tplot(t)=t;
   y(t)=y(t-1)+mu_step+sqrt(var_step)*randn;
end
tlabel=zeros(1,10);
ylabell=zeros(1,10);
err=zeros(1,10);

j=1;
for j=1:10
```

```
    for t=1:1000
        if t/100==j
            tlabel(j)=t-99*rand;
            j=j+1;
        end
    end
end
tlabel=round(tlabel);
for j=1:10
    n(1,j)=57*rand;
end
n=round(n)+3*ones(1,10);

j=1;
for t=1:1000
    if t==tlabel(j)
        individuals=sqrt(Vp)*randn(n(j),1);
        Mean(1,j)=mean(individuals);
        ytrue(j)=y(t);
        ylabell(j)=y(t)+Mean(1,j);
        err(j)=sqrt(Vp/n(j));
        if j==10 break
        end
        j=j+1;
    end
end

%% GLS on true y data

Cov0=zeros(N,N);
for i=1:N
    for j=1:N
        Cov0(i,j)=min(tlabel(i),tlabel(j));
    end
end

Cov1=Cov0*var_step;
```

```
v=err.^2;
V=diag(v);

X=[ones(10,1) tlabel'];
bls=inv(X'*inv(Cov1+V)*X)*X'*inv(Cov1+V)*ytrue';
a_GLS=bls(1);
b_GLS(m)=bls(2);
for i=1:10
   yhat_GLS(i)=a_GLS+b_GLS(m)*tlabel(i);
end
% WMSE_GLS(m)=(ytrue-yhat_GLS)*inv(Cov1+V)*(ytrue-yhat_GLS)'/trace(inv(Cov1+V));

%% Hunt's GRW

n_samples=ones(1,10);
var_samples=err.^2;

c=1;
Tsample=c*tlabel;

for i=1:9
   dT(i)=-Tsample(i)+Tsample(i+1);
   dX(i)=ylabell(i+1)-ylabell(i);
   nA(i)=n_samples(i);
   nD(i)=n_samples(i+1);
   varA(i)=var_samples(i);
   varD(i)=var_samples(i+1);
end

% Contraints
mustep_min=-100; mustep_max=100;
varstep_min=0; varstep_max=100;
par_lb=[mustep_min varstep_min];
par_ub=[mustep_max varstep_max];

% fmincon
par_guess=[0 0];
Aineq=[]; Bineq=[]; Aeq=[]; Beq=[];
```

```
fun_objective_handle=...
   @(par)fun_objective(par,dT,dX,varA,nA,varD,nD);
[par_opt,fval,exitflag,output,lambda,grad,hessian] =...
fmincon(fun_objective_handle,par_guess,Aineq,Bineq,Aeq,Beq,par_lb,par_ub);

mustep(m)=par_opt(1);
varstep(m)=par_opt(2);

if varstep(m)<0
   count=count+1;
end

X=ones(10,1);
b_GRW(m)=c*mustep(m);
a_GRW(m)=inv(X'*inv(V)*X)*X'*inv(V)*(ylabell'-b_GRW(m)*tlabel');
for j=1:10
   yhat_GRW(j)=a_GRW(m)+b_GRW(m)*tlabel(j);
end

%% WLS

X=[ones(10,1) tlabel'];
bls=inv(X'*inv(V)*X)*X'*inv(V)*ytrue';
a_WLS=bls(1);
b_WLS(m)=bls(2);
for i=1:10
   yhat_WLS(i)=a_WLS+b_WLS(m)*tlabel(i);
end

end

count

figure(1)
subplot(3,2,1)
histogram(b_GLS,40,'BinWidth',0.005,'FaceColor','b')
ax = findobj(subplot(3,2,1),'Type','Axes');
for i = 1:length(ax)
```

```
    ylim(ax(i),[0 120]);
    xlim(ax(i),[0.04 0.16]);
end
title('Hunt´s AD model')
xlabel('Prediction slope b_G_L_S')

subplot(3,2,3)
histogram(b_GRW-b_GLS,40,'BinWidth',0.005,'FaceColor','m')
ax = findobj(subplot(3,2,3),'Type','Axes');
for i = 1:length(ax)
    ylim(ax(i),[0 400]);
    xlim(ax(i),[-0.06 0.06]);
end
xlabel('Prediction slope error, b_G_R_W - b_G_L_S')

subplot(3,2,5)
histogram(b_WLS-b_GLS,40,'BinWidth',0.005,'FaceColor','m')
ax = findobj(subplot(3,2,5),'Type','Axes');
for i = 1:length(ax)
    ylim(ax(i),[0 400]);
    xlim(ax(i),[-0.06 0.06]);
end
xlabel('Prediction slope error, b_W_L_S - b_G_L_S')

%% Parameter search
function f = fun_objective(par,dT,dX,varA,nA,varD,nD)
mustep=par(1);
varstep=par(2);

for i=1:9
    varterm(i)=dT(i)*varstep+varA(i)/nA(i)+varD(i)/nD(i);
    Li(i)=-log(2*pi)/2-log(varterm(i))/2-(dX(i)-dT(i)*mustep)^2/(2*varterm(i));
end

L=sum(Li);

f=-L;
```

```
end
```

#### Figure 1, right panels, and Figure 2

```
clear

mu_step=0.1;
var_step=0.4;
Vp=400;
count=0;
y0=0;
M=1000
N=10

for m=1:M
%% Generate data
y=y0*zeros(1,1000);
for t=2:1000
   tplot(t)=t;
   y(t)=y(t-1)+mu_step+sqrt(var_step)*randn;
end
tlabel=zeros(1,10);
ylabell=zeros(1,10);
err=zeros(1,10);

j=1;
for j=1:10
   for t=1:1000
      if t/100==j
         tlabel(j)=t-99*rand;
         j=j+1;
      end
   end
end
tlabel=round(tlabel);
for j=1:10
   n(1,j)=57*rand;
end
```

```
n=round(n)+3*ones(1,10);


j=1;
for t=1:1000
   if t==tlabel(j)
      individuals=sqrt(Vp)*randn(n(j),1);
      Mean(1,j)=mean(individuals);
      ytrue(j)=y(t);
      ylabell(j)=y(t)+Mean(1,j);
      err(j)=sqrt(Vp/n(j));
      if j==10 break
      end
      j=j+1;
   end
end

%% GLS

Cov0=zeros(N,N);
for i=1:N
   for j=1:N
      Cov0(i,j)=min(tlabel(i),tlabel(j));
   end
end

Cov1=Cov0*var_step;

v=err.^2;
V=diag(v);

X=[ones(10,1) tlabel'];
bls=inv(X'*inv(Cov1+V)*X)*X'*inv(Cov1+V)*ytrue';
a_GLS=bls(1);
b_GLS(m)=bls(2);
for i=1:10
   yhat_GLS(i)=a_GLS+b_GLS(m)*tlabel(i);
end
```

```
WMSE_GLS(m)=(ytrue-yhat_GLS)*inv(Cov1+V)*(ytrue-yhat_GLS)'/trace(inv(Cov1+V));


%% GRW


n_samples=ones(1,N);

var_samples=v;


c=1;

Tsample=c*tlabel;


for i=1:N-1

   dT(i)=Tsample(i+1)-Tsample(i);

   dX(i)=ylabell(i+1)-ylabell(i);

   nA(i)=n_samples(i);

   nD(i)=n_samples(i+1);

   varA(i)=var_samples(i);

   varD(i)=var_samples(i+1);

end


mustep_min=-100; mustep_max=100;

varstep_min=0; varstep_max=100;

% varstep_min=-0.000001; varstep_max=0.000001;

par_lb=[mustep_min varstep_min];

par_ub=[mustep_max varstep_max];


% fmincon

par_guess=[0 0];

Aineq=[]; Bineq=[]; Aeq=[]; Beq=[];

fun_objective_handle=...

   @(par)fun_objective(par,dT,dX,varA,nA,varD,nD,v,tlabel,N);

[par_opt,fval,exitflag,output,lambda,grad,hessian] =...

fmincon(fun_objective_handle,par_guess,Aineq,Bineq,Aeq,Beq,par_lb,par_ub);


mustep(m)=par_opt(1);

varstep(m)=par_opt(2);


var_main=varstep(m);
```

```
for i=1:N-1
   H(i)=(dX(i)-dT(i)*mustep(m));
   D_main(i)=v(i)+v(i+1)+dT(i)*var_main;
end

for i=1:N-2
   D_side(i)=-v(i+1);
end

Sigma_1=diag(D_main);

Sigma_2=[zeros(1,N-1)
      diag(D_side) zeros(N-2,1)];

Sigma_3=[zeros(N-2,1) diag(D_side)
       zeros(1,N-1)];

Sigma=Sigma_1+Sigma_2+Sigma_3;

L=-(N-1)*log(2*pi)/2-log(det(Sigma))/2-H*inv(Sigma)*H'/2;

NAIC=N-1;
k=2;

AICc_GRW(m)=2*k-2*L+2*k*(k+1)/(NAIC-k-1);

X=ones(N,1);
b_GRW(m)=mustep(m);
a_GRW(m)=inv(X'*inv(V)*X)*X'*inv(V)*(ylabell'-b_GRW(m)*tlabel');
for j=1:N
   yhat_GRW(j)=a_GRW(m)+b_GRW(m)*tlabel(j);
end

%% WLS

X=[ones(10,1) tlabel'];
bls=inv(X'*inv(V)*X)*X'*inv(V)*ytrue';
a_WLS(m)=bls(1);
```

```
b_WLS(m)=bls(2);
for i=1:10
   yhat_WLS(i)=a_WLS(m)+b_WLS(m)*tlabel(i);
end

end

figure(1)
subplot(3,2,2)
histogram(b_GLS,40,'BinWidth',0.005,'FaceColor','b')
ax = findobj(subplot(3,2,2),'Type','Axes');
for i = 1:length(ax)
   ylim(ax(i),[0 120]);
   xlim(ax(i),[0.04 0.16]);
end
title('Corrected AD model')
xlabel('Prediction slope b_G_L_S')

subplot(3,2,4)
histogram(b_GRW-b_GLS,40,'BinWidth',0.005,'FaceColor','m')
ax = findobj(subplot(3,2,4),'Type','Axes');
for i = 1:length(ax)
   ylim(ax(i),[0 400]);
   xlim(ax(i),[-0.06 0.06]);
end
xlabel('Prediction slope error, b_G_R_W - b_G_L_S')

subplot(3,2,6)
histogram(b_WLS-b_GLS,40,'BinWidth',0.005,'FaceColor','m')
ax = findobj(subplot(3,2,6),'Type','Axes');
for i = 1:length(ax)
   ylim(ax(i),[0 400]);
   xlim(ax(i),[-0.06 0.06]);
end
xlabel('Prediction slope error, b_W_L_S - b_G_L_S')

figure(2)
plot(tplot,y), hold on
```

```
errorbar(tlabel,ylabell,err,'bo','LineWidth',1.0), hold on
plot(tlabel,yhat_GLS,'g','LineWidth',1.0)
plot(tlabel,yhat_WLS,'--m','LineWidth',1.0)
plot(tlabel,yhat_GRW,'m','LineWidth',1.0)
% plot(tlabel,ytrue,'ks')
% plot(tlabel,ytrue,'ko')

hold off, grid
xlabel('time step')
ylabel('trait value')
title('V_p = 400')
axis([0 1000 0 100])

%% Parameter search
function f = fun_objective(par,dT,dX,varA,nA,varD,nD,v,tlabel,N)
mustep=par(1);
varstep=par(2);

var_main=varstep;

for i=1:N-1
   H(i)=(dX(i)-dT(i)*mustep);
   D_main(i)=v(i)+v(i+1)+dT(i)*var_main;
end

for i=1:N-2
   D_side(i)=-v(i+1);
end

Sigma_1=diag(D_main);

Sigma_2=[zeros(1,N-1)
      diag(D_side) zeros(N-2,1)];

Sigma_3=[zeros(N-2,1) diag(D_side)
       zeros(1,N-1)];

Sigma=Sigma_1+Sigma_2+Sigma_3;
```

```
L=-(N-1)*log(2*pi)/2-log(det(Sigma))/2-H*inv(Sigma)*H'/2;

f=-L;

end
```

#### Figure 3, left panel

```
clear

N=10

s2=467;

%% Sample data
Data=[
3.33 1,962 2.54 13 707
2.60 3,427 2.23 4 744
2.40 1,253 2.15 21 742
2.19 1,253 2.07 23 743
1.90 1,253 1.96 25 745
1.68 1,253 1.88 22 763
0.89 2,873 1.55 28 812
0.14 2,873 1.19 24 797
0.12 3,427 1.18 3 793
0.00 2,781 1.12 31 812];

tlabel=-Data(:,1)';
y=Data(:,6)';
ylabell=log(Data(:,6))';
n=Data(:,5)';

for i=1:N
   err(i)=sqrt(s2/n(i))*ylabell(i)/y(i);
end

w=err.^-2;
```

```
W=diag(w);
v=err.^2;
V=diag(v);

%% WLS model

X=[ones(10,1) tlabel'];
bls=inv(X'*W*X)*X'*W*ylabell';
a_WLS=bls(1);
b_WLS=bls(2);
for i=1:10
   yhat_WLS(1,i)=a_WLS+b_WLS*tlabel(i);
end

%% GRW model

n_samples=ones(1,N);
var_samples=err.^2;

c=1;
Tsample=c*tlabel;

for i=1:N-1
   dT(i)=-Tsample(i)+Tsample(i+1);
   dX(i)=ylabell(i+1)-ylabell(i);
   nA(i)=n_samples(i);
   nD(i)=n_samples(i+1);
   varA(i)=var_samples(i);
   varD(i)=var_samples(i+1);
end

% Contraints
mustep_min=-1; mustep_max=1;
varstep_min=0; varstep_max=1;

par_lb=[mustep_min varstep_min];
par_ub=[mustep_max varstep_max];
```

```
par_guess=[0 0];
Aineq=[]; Bineq=[]; Aeq=[]; Beq=[];
fun_objective_handle=...
   @(par)fun_objective(par,dT,dX,varA,nA,varD,nD,v,N);
[par_opt,fval,exitflag,output,lambda,grad,hessian] =...
fmincon(fun_objective_handle,par_guess,Aineq,Bineq,Aeq,Beq,par_lb,par_ub);

par_opt;
varstep=par_opt(2);
b_GRW=par_opt(1);

var_main=varstep;

for i=1:N-1
   H(i)=(dX(i)-dT(i)*b_GRW);
   D_main(i)=v(i)+v(i+1)+dT(i)*var_main;
end

for i=1:N-2
   D_side(i)=-v(i+1);
end

Sigma_1=diag(D_main);

Sigma_2=[zeros(1,N-1)
      diag(D_side) zeros(N-2,1)];

Sigma_3=[zeros(N-2,1) diag(D_side)
       zeros(1,N-1)];

Sigma=Sigma_1+Sigma_2+Sigma_3;

L=-(N-1)*log(2*pi)/2-log(det(Sigma))/2-H*inv(Sigma)*H'/2;

NAIC=N-1;
k=2;

AICc_GRW=2*k-2*L+2*k*(k+1)/(NAIC-k-1);
```

```
X=ones(10,1);

a_GRW=inv(X'*W*X)*X'*W*(ylabell'-b_GRW*tlabel');

for i=1:10

   yhat_GRW(i)=a_GRW+b_GRW*tlabel(i);

end


varstep

b_GRW

b_WLS

AICc_GRW


figure(3)

subplot(1,2,1)

errorbar(tlabel,ylabell,err,'bo','LineWidth',1.0), hold on

plot(tlabel,yhat_WLS,'b','LineWidth',1.0)

plot(tlabel,yhat_GRW,'--m','LineWidth',1.0), hold off, grid

axis([-3.4 0.1 6.5 6.8])

xlabel('Million years')

ylabel('log mean body size')

title('Ostracod 1 case')


%% Parameter search

function f = fun_objective(par,dT,dX,varA,nA,varD,nD,v,N)

mustep=par(1);

varstep=par(2);


var_main=varstep;


for i=1:N-1

   H(i)=(dX(i)-dT(i)*mustep);

   D_main(i)=v(i)+v(i+1)+dT(i)*var_main;

end


for i=1:N-2

   D_side(i)=-v(i+1);

end
```

```
Sigma_1=diag(D_main);

Sigma_2=[zeros(1,N-1)
    diag(D_side) zeros(N-2,1)];

Sigma_3=[zeros(N-2,1) diag(D_side)
     zeros(1,N-1)];

Sigma=Sigma_1+Sigma_2+Sigma_3;
% Sigma=Sigma_1;

L=-(N-1)*log(2*pi)/2-log(det(Sigma))/2-H*inv(Sigma)*H'/2;

f=-L;

end
```
